## Supplementary material for "The impact of HTLV-1 expression on the 3D structure and expression of host chromatin": All supplemental information

#### S1 Fig

##### Clone 11.63

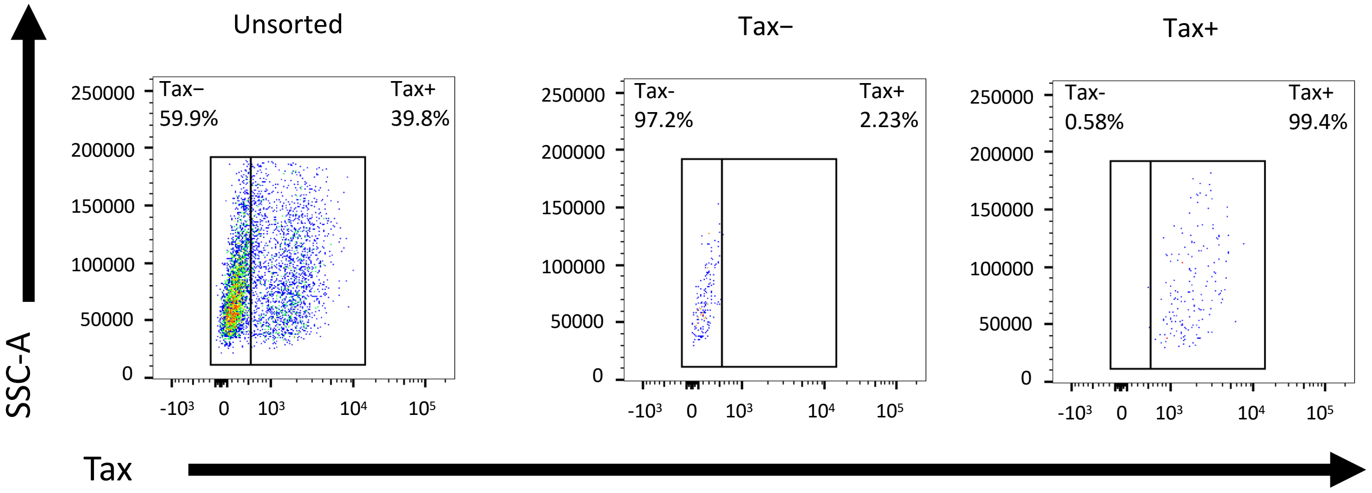

#### S2 Fig

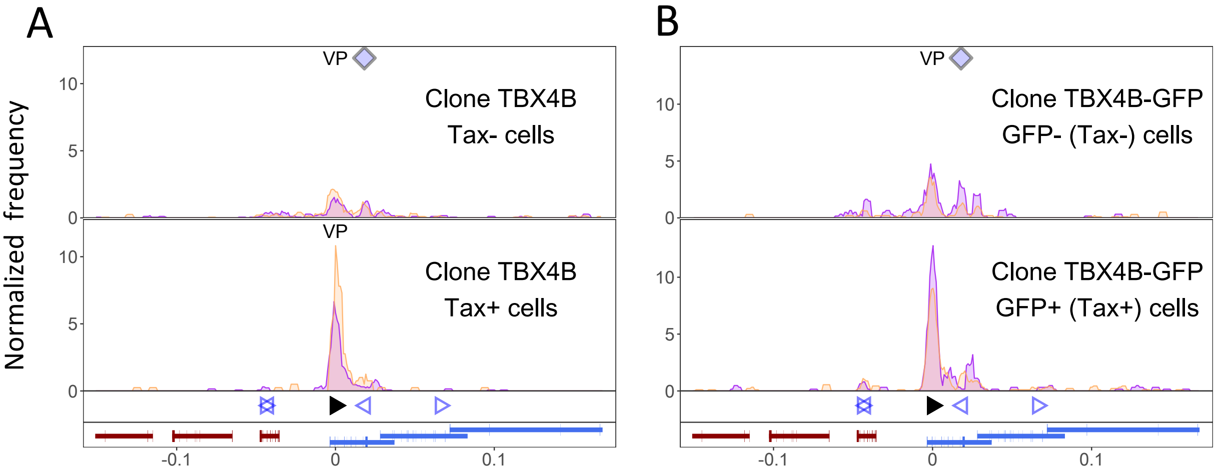

A
B
VP
VP
Clone TBX4B
Tax- cells
Clone TBX4B
Tax+ cells
Clone TBX4B-GFP
GFP- (Tax-) cells
Clone TBX4B-GFP
GFP+ (Tax+) cells
Normalized frequency
10
5
0
-0.1
0
0.1

#### S3 Fig

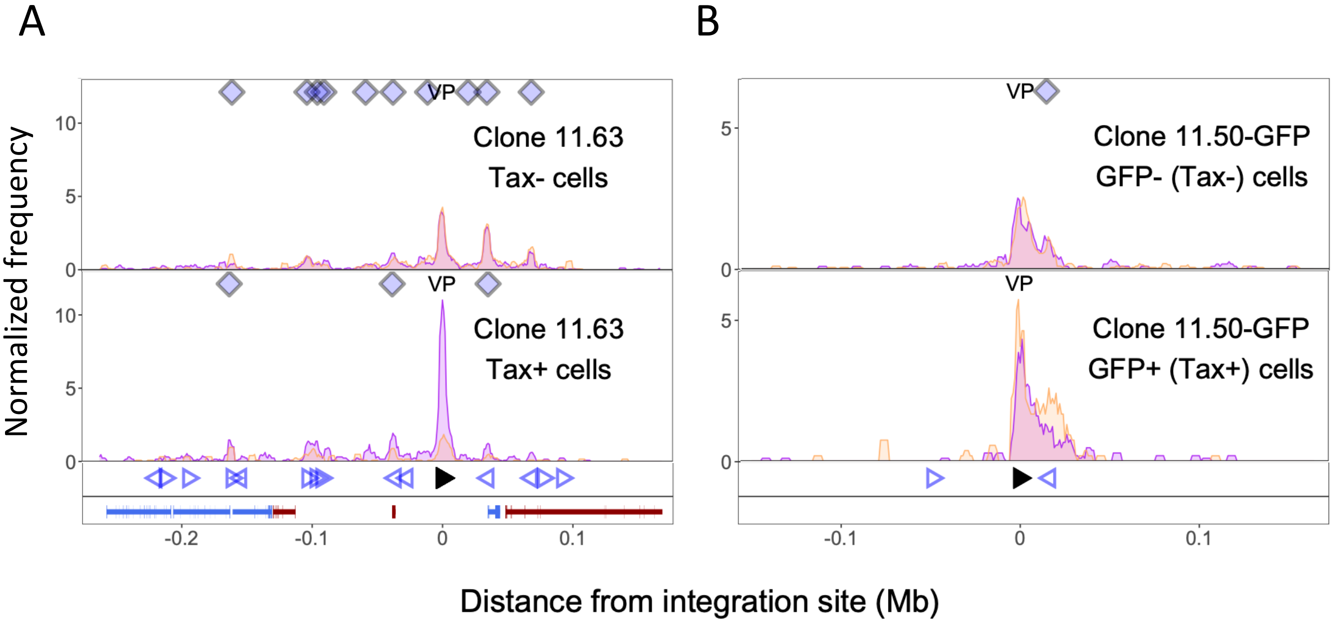

A
B
Normalized frequency
Clone 11.63
Tax- cells
Clone 11.63
Tax+ cells
Clone 11.50-GFP
GFP- (Tax-) cells
Clone 11.50-GFP
GFP+ (Tax+) cells
VP
10
5
0
-0.2
-0.1
0
0.1
Distance from integration site (Mb)

### S4 Fig

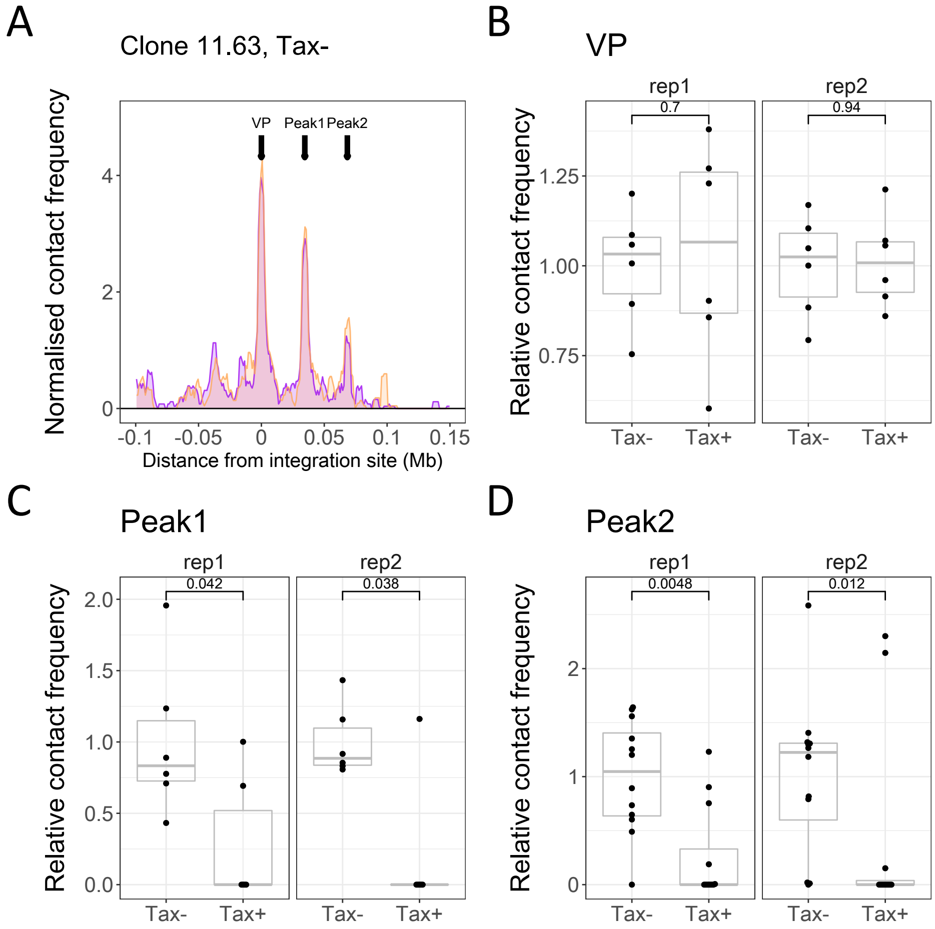
A
Clone 11.63, Tax-
VP
Peak1
Peak2
Normalised contact frequency
Distance from integration site (Mb)
B
VP
rep1
rep2
0.7
0.94
Relative contact frequency
Tax-
Tax+
C
Peak1
0.042
0.038
D
Peak2
0.0048
0.012

### S5 Fig

#### A Clone 11.63

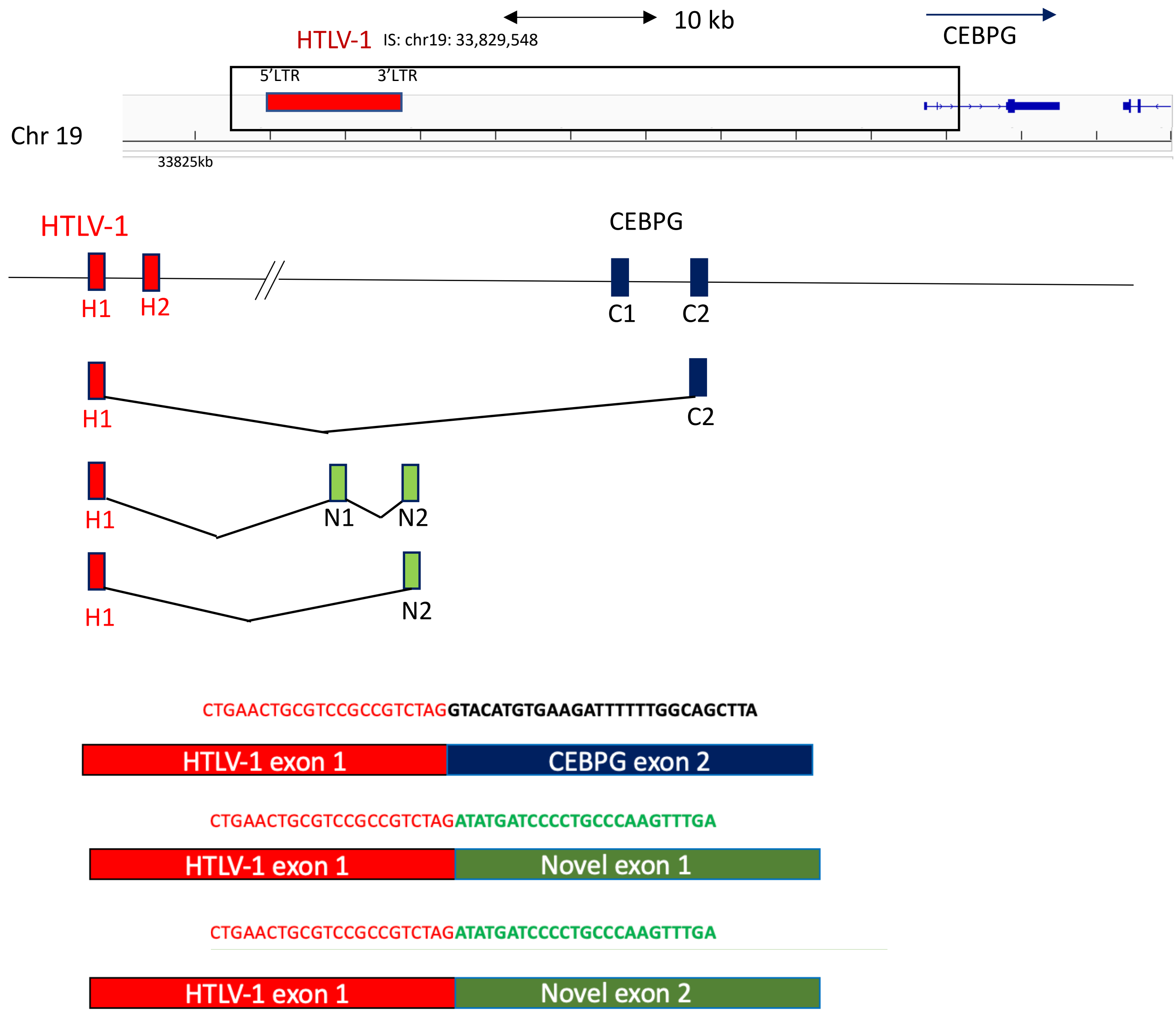

B Clone Timer-3.60

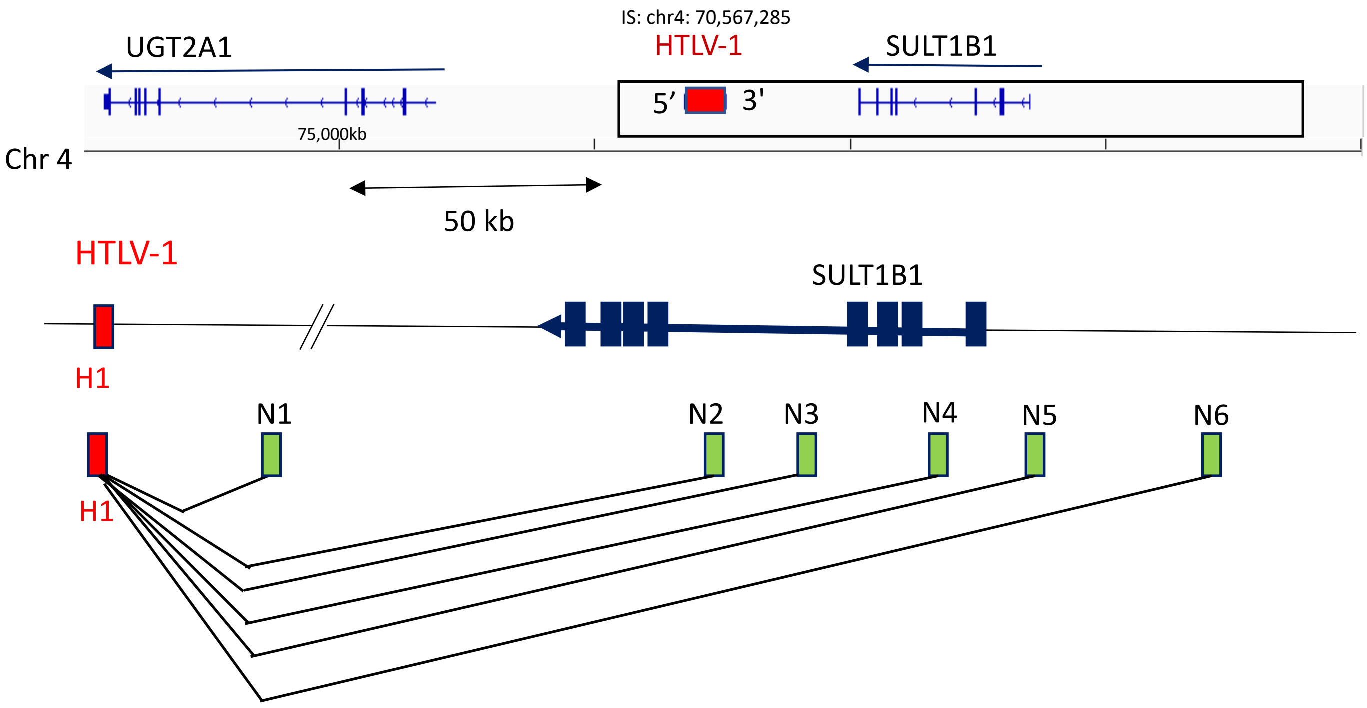

### S6 Fig

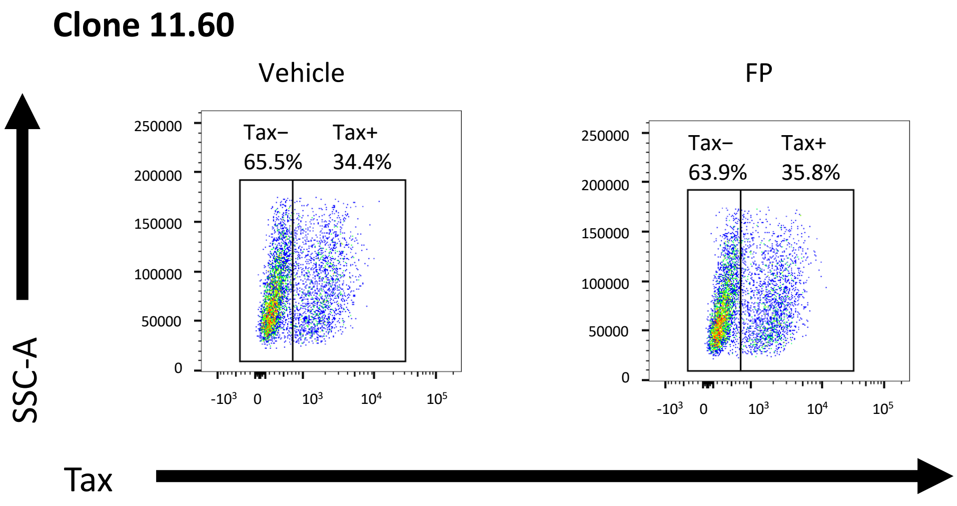

Clone 11.60
Vehicle
FP
Tax−
65.5%
Tax+
34.4%
Tax−
63.9%
Tax+
35.8%
250000
200000
150000
100000
50000
0
-10³
0
10³
10⁴
10⁵
SSC-A
Tax

**S7 Fig**

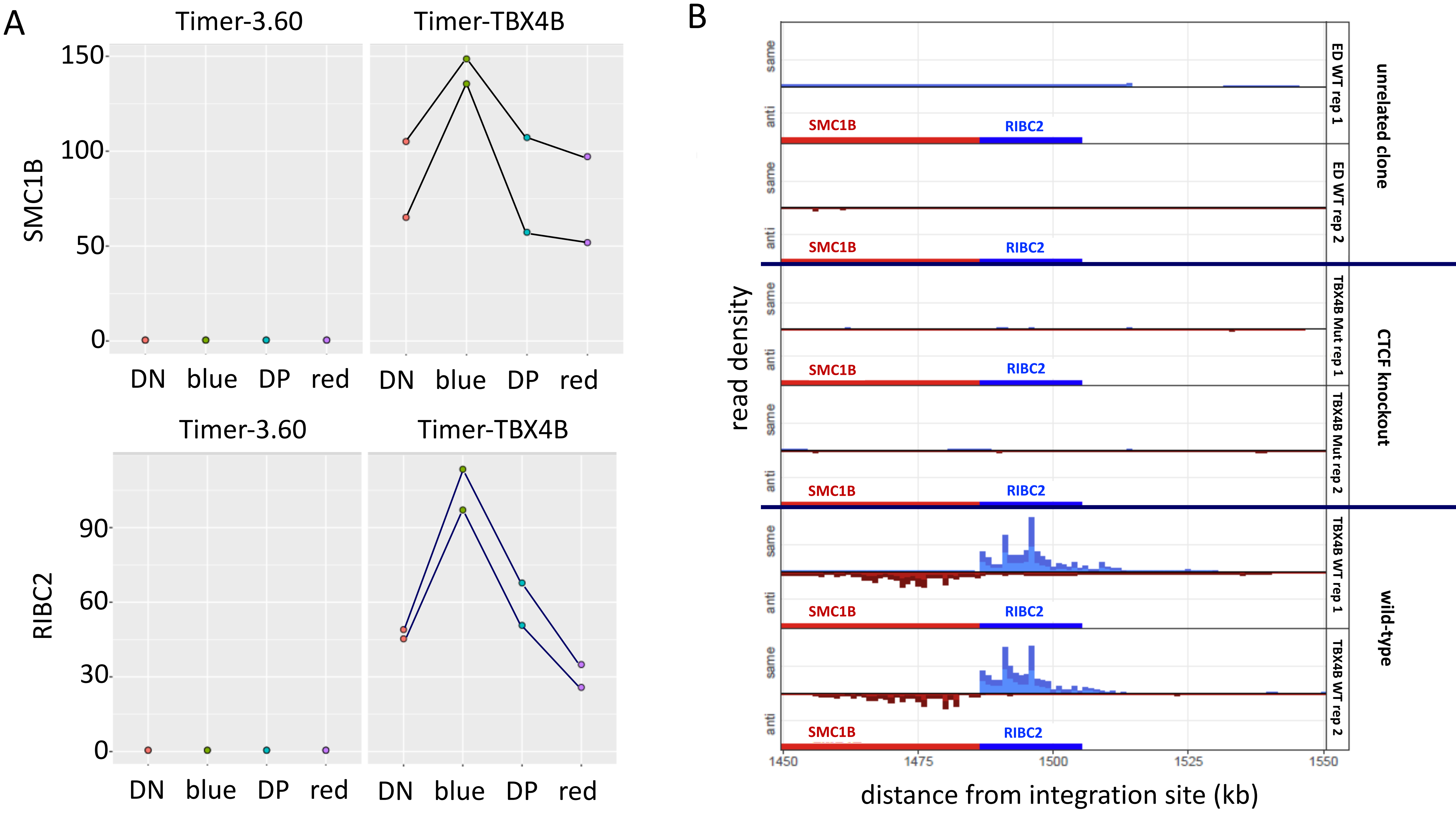
A
Timer-3.60
Timer-TBX4B
SMC1B
150
100
50
0
DN
blue
DP
red
Timer-3.60
Timer-TBX4B
RIBC2
90
60
30
0
DN
blue
DP
red
B
read density
same
anti
SMC1B
RIBC2
ED WT rep 1
ED WT rep 2
unrelated clone
TBX4B Mut rep 1
TBX4B Mut rep 2
CTCF knockout
TBX4B WT rep 1
TBX4B WT rep 2
wild-type
1450
1475
1500
1525
1550
distance from integration site (kb)

**S8 Fig**

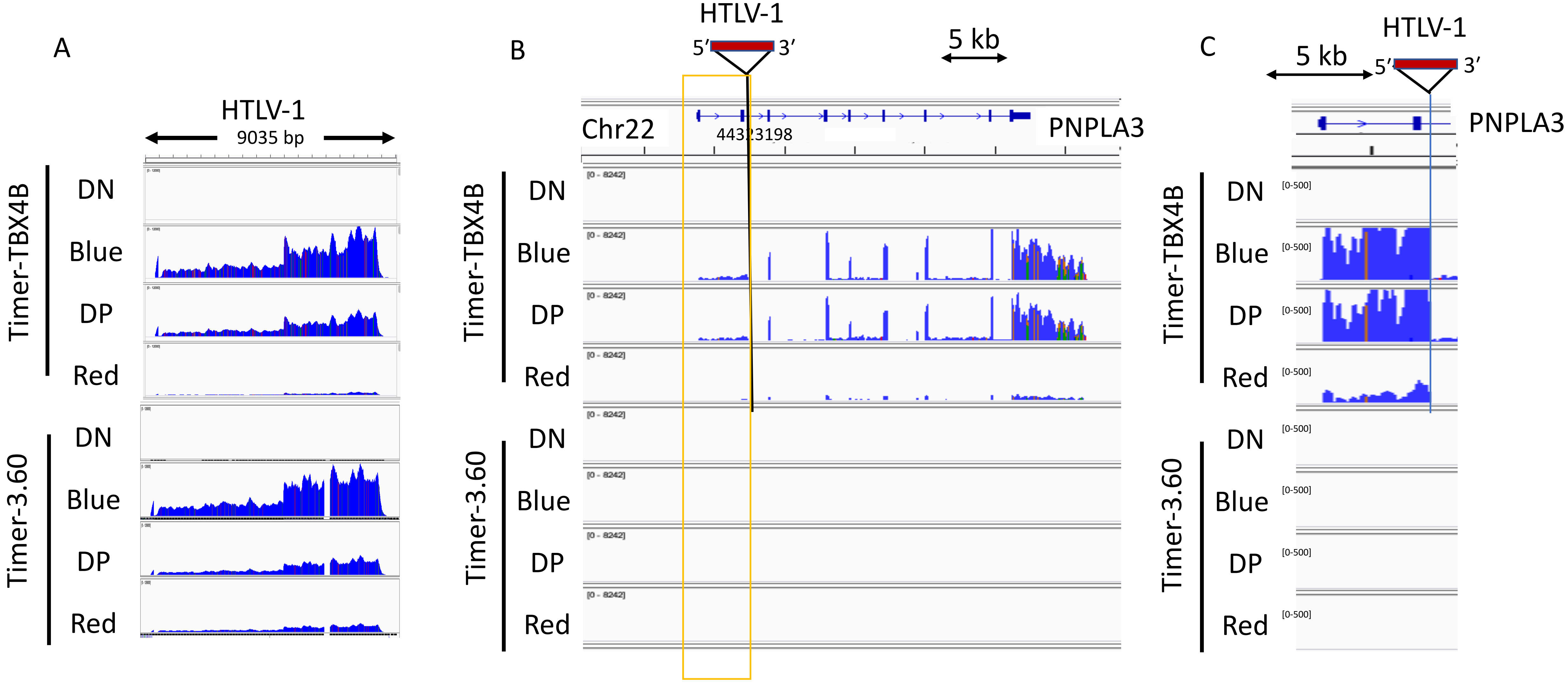
A
HTLV-1
9035 bp
Timer-TBX4B
DN
Blue
DP
Red
Timer-3.60
DN
Blue
DP
Red
B
HTLV-1
5′
3′
5 kb
Chr22
44323198
PNPLA3
Timer-TBX4B
DN
Blue
DP
Red
Timer-3.60
DN
Blue
DP
Red
[0 - 8242]
C
HTLV-1
5 kb
5′
3′
PNPLA3
Timer-TBX4B
DN
Blue
DP
Red
Timer-3.60
DN
Blue
DP
Red
[0-500]

**S1 Table**

**Primers and probes used**

| Experiment | Target | Primer and probe names | Sequence (5'-3') |
|---|---|---|---|
| 3C-qPCR | Peak 1 | HTLV1_7200_7220 | TAATAGCCCGTCCACCAATTC |
| | | Peak1_Rev | AACCAGCCTGGACAACATAG |
| | | Peak1_Probe | TCCGGGCATGTGACATCAATGCTA |
| | Peak 2 | HTLV1_7173 | CTTGTCCTTTAACTCTTCCTCCA |
| | | Peak2_Rev | CAGTGTTGTCTCAGGACGATG |
| | | Peak2_Probe | AATAGCCCGTCCACCAATTCCTCC |
| | Control | HTLV1_7200_7220 | TAATAGCCCGTCCACCAATTC |
| | | HTLV1_6319_6301 | TCTGTTCTGGGCAGCATAC |
| | | VP_probe | CCTCCGGGCATGAGGTGGACAAAGATA |
| | Internal control | HTLV1_7118_7136 | CGCGGCTTTCCTCTTCTAA |
| | | HTLV1_7218_7198 | ATTGGTGGACGGGCTATTATC |
| | | C_Probe | AAGCACAGCTTCCTCCTCCTCCTT |
| RT-qPCR | 18S rRNA | 18S_Fwd | GTAACCCGTTGAACCCCATT |
| | | 18S_Rev | CCATCCAATCGGTAGTAGCG |
| | Tax | Tax_Fwd | CCGGCGCTGCTCTCATCCCGGT |
| | | Tax_Rev | GGCCGAACATAGTCCCCCAGAG |
| | 11.63 IS plus 188-380bp | PositionA_Fwd | CCACCATGCTTGGCTACTT |
| | | PositionA_Rev | TTCTGCCCTTAACTCTGGAATG |
| | 11.63 IS plus 535-648bp | PositionB_Fwd | GGACTTTACCAGAAGCAACCT |
| | | PositionB_Rev | TTCCCTTCCCTACCTCACTT |
| | 11.63 IS plus 3198 to 3304bp | PositionC_Fwd | GCGGAAAAGCCCAAGTTTGA |
| | | PositionC_Rev | CTCATTCAACTTTAGCACCGG |
